# Squeakuences: a portable tool for formatting ‘squeaky-clean’ sequences to eliminate bioinformatic software incompatibilities

**DOI:** 10.1101/2024.11.01.621607

**Authors:** Linnea E Lane, Ashley N Doerfler, Luna A L’Argent, Emily Touchette, Bronson Mills, Jessika Bryant, Savannah N Roller, Evan S Forsythe

## Abstract

Computational analysis of biological sequences is the cornerstone of modern bioinformatics research. Complex processing and interpretation of data often entails multi-step workflows. The specific requirements and limitations of individual applications can require laborious reformatting and piecemeal ‘data-wrangling’ to produce a satisfactory input for each step in a pipeline. We present *Squeakuences*, a command line tool developed to simplify and automate FASTA file preparation for applications such as phylogenetics, gene annotation, and genome analysis. Implemented in a lightweight Python script, *Squeakuences* identifies and removes potentially problematic elements in sequence identifiers, such as non-alphanumeric characters, white space, and excessive character count. *Squeakuences* outputs a new clean version of the sequence file for analysis alongside metadata files to track changes. The user can customize *Squeakuences*’ behavior using optional arguments to meet individual processing and formatting requirements. We tested the performance of *Squeakuences* on molecular data from the human reference genome and found that runtime correlates with the number of sequences processed but not with file size. We expect *Squeakuences* to save time and manual effort when analyzing sequence data. *Squeakuences* code is freely available at https://github.com/EvanForsythe/Squeakuences.

## Introduction

FASTA files are a popular format for storing sequence information and are ubiquitous in molecular and evolutionary biology. Bioinformatic research workflows are often implemented as multistep ‘pipelines’ that incorporate several software programs. While most bioinformatic programs accept FASTA-formatted files, each program tends to have its own set of formatting requirements, meaning variations in sequence identifier (seq ID) contents such as whitespace, punctuation, and special characters can raise errors or produce unexpected behavior, inhibiting the analysis process. This issue necessitates tedious manual editing of FASTA files to remove invalid characters and repeated preprocessing at each analysis step. Web-based (Foley et al. 2019) and graphical-user-interface-based (Kearse et al. 2012) tools provide batch reformatting functions; however, such programs are not easily incorporated into command-line based workflows, including those that are implemented in high-performance distributed computing environments.

Here, we introduce *Squeakuences*, a lightweight python-based tool for simple and flexible FASTA file reformatting. *Squeakuences* is designed to systematically address all possible issues that could arise from seq ID formatting, freeing researchers from the need to anticipate and manually address all possible formatting obstacles. We tested *Squeakuences* on protein, RNA, and DNA sequences and found that the tool successfully cleans each of these file types. *Squeakuences* will facilitate bioinformatic analyses and help make computational analyses accessible to researchers with any level of prior computer science experience.

## Methods

*Squeakuences* utilizes regular expressions and built-in functions to identify and resolve problematic content in seq ID strings. In addition to replacing problematic characters, *Squeakuences* also intelligently trims seq IDs that surpass a user-defined limit, such that whole words are retained in sequence descriptions, thus minimizing information loss. Additionally, *Squeakuences* resolves identical seq IDs within a FASTA file and provides optional functionality to prepend species IDs to seq ID strings, thus facilitating comparative analyses that merge sequences from multiple species. *Squeakuences* does not require additional software dependencies, so it can be run in the user’s preferred *Python 3* environment. *Squeakuences* can process a single file or batch process multiple files. Following several cleaning and formatting steps (Fig. 1), *Squeakuences* writes a new FASTA file containing the cleaned seq IDs and their corresponding unaltered sequence data. Changes to seq IDs are tracked in a tab-separated metadata file for record keeping and back-compatibility.

**Figure 1:**
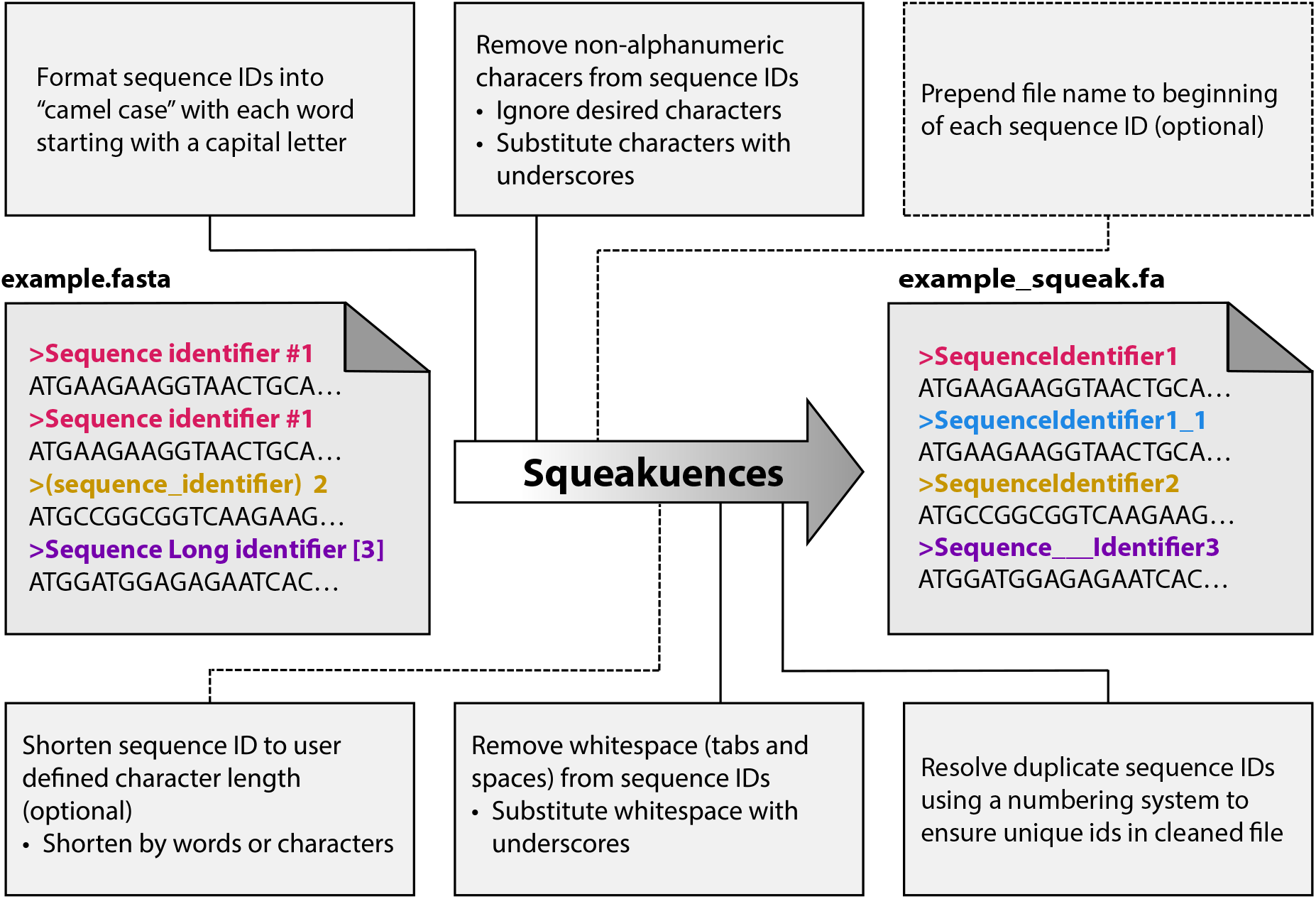
Graphical abstract of Squeakuences workflow. Major processes implemented in *Squeakuences*. Analytical steps are shown in the order they are performed. Dotted line indicates optional steps, and sub-bullets indicate customization arguments.

*Squeakuences* is run with the minimal command, `‘python3 squeakuences -i example.fa -o example_out_dir`/’, where ‘-i’ and ‘-o’ are required arguments that specify the input file or directory and output directory, respectively. We tested *Squeakuences* (v1.1.0) using python (v3.12) on *macOS* and *Linux* systems and made use of the optional ‘--log’ argument to track performance across different file types.

## Results

To understand the performance of *Squeakuences* on a variety of FASTA files, we processed RNA, genomic DNA, and protein FASTA files for the human reference genome, HG38 (NCBI accession: GCF_000001405.40) (Fig. 2). Files varied in the number and length of seq IDs and the length of associated sequence data, with some containing only a few long sequences while others contain a large number of short sequences (Fig. 2A). We found that the runtime of *Squeakuences* exhibits a general positive linear correlation with the number of seq IDs cleaned (Fig. 2B) but not with overall file size (Fig. 2E), meaning that the number of seq IDs processed is the primary driver of *Squeakuences* runtime. We also tracked peak memory usage of runs and found that memory usage is tightly correlated to file size, meaning memory requirements will scale with overall file size, although no evidence of memory-related performance reduction was detected in our tests. Our tests show that *Squeakuences* completes in less than seven minutes and requires less than 6 GB of memory, even on the largest datasets. This efficiency makes *Squeakuences* lightweight enough to run on both distributed computing systems and standard laptops.

**Figure 2:**
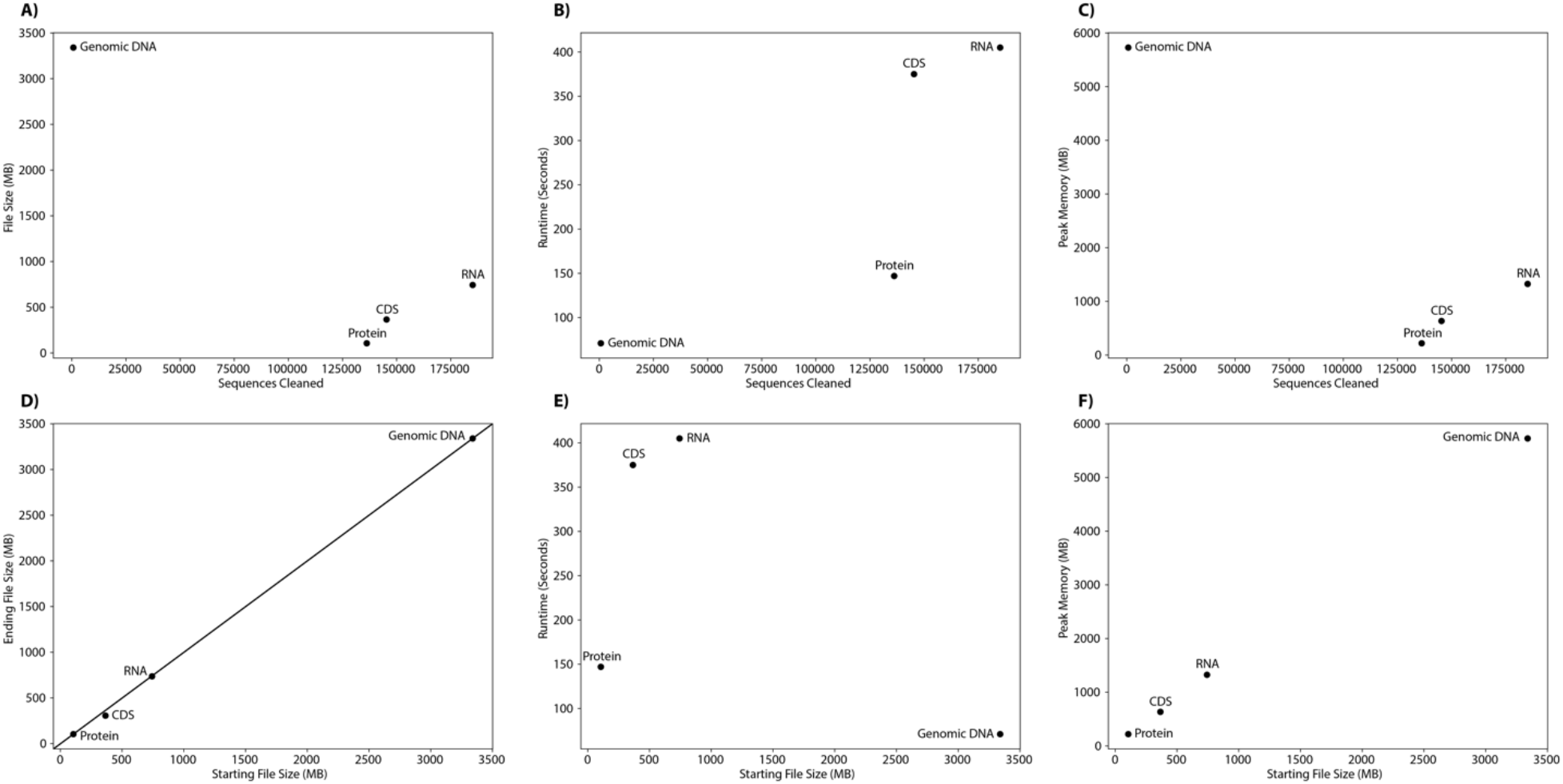
Squeakuences performance on a variety of file types. FASTA files containing hg38 genomic data were processed by Squeakuences. (A-C) Scatterplots comparing the number of sequences cleaned to the starting file size (A), runtime (B), and peak memory usage (C). (D-F) Scatterplots comparing starting file size to ending file size (D), runtime (E), and peak memory used (F). Diagonal line in panel D indicates one-to-one line.

## Discussion

Squeakuences was created to address common stumbling blocks experienced by bioinformatics researchers. The current targets for seq ID cleaning (white space, non-alphanumeric characters, exceedingly long strings, duplicate IDs) are based on issues experienced by the authors in their own real-world bioinformatic applications; however, it is likely that additional unforeseen formatting issues are not solved by the current version of *Squeakuences* and will come to light as more users utilize *Squeakuences*. We plan to use the *Squeakuences* GitHub page as an open forum to continually gather examples of formatting issues and to consistently update *Squeakuences*. Taken together, *Squeakuences* will be a valuable tool for resolving formatting issues in order to bring efficiency and peace-of-mind to bioinformatic research.

## Acknowledgements

The testing and benchmarking stages of this work were performed by student researchers as part of a course-based undergraduate research experience (CURE) in the Applied Bioinformatics (BB485) course at OSU-Cascades. This work was supported by a National Science Foundation (NSF) grant (IOS-2114641).

## Citations

Foley G, Sützl L, D’Cunha SA, Gillam EMJ, and Bodén M. SeqScrub: A web tool for automatic cleaning and annotation of FASTA file headers for bioinformatic applications. Biotechniques. 2019:67(2):50–54. 10.2144/btn-2018-0188

Kearse M, Moir R, Wilson A, Stones-Havas S, Cheung M, Sturrock S, Buxton S, Cooper A, Markowitz S, Duran C, et al. Geneious Basic: An integrated and extendable desktop software platform for the organization and analysis of sequence data. Bioinformatics. 2012:28(12):1647–1649. 10.1093/bioinformatics/bts199

